# Lie Detectors for Face Recognition

**DOI:** 10.1101/2025.06.22.660938

**Authors:** Marsha Meytlis

## Abstract

Lie detection is important for government law enforcement. Current lie detection methods such as the polygraph test have been found to be unreliable (Meijer *et al*. 2017). New lie detection technology is currently arising that is based on fMRI; however, single subject tests have only been successful in detecting lies 88% of the time (Langleben *et al*., 2005; Wild, 2005). One of the main problems with most fMRI-based approaches is that they assume that various acts of deception involve common brain regions, (Ganis *et al*., 2003). In this work I propose a much more accurate fMRI lie detection method that does not make this assumption and is domain based. In my investigation, rather than trying to localize brain regions that are indicative of lying in general, I localize brain regions that indicate lying specifically about face recognition. In criminal investigations one frequently needs to establish familiar relations between a suspect, victim and/or witness. This type of information can be used as circumstantial evidence in a crime. In this work I propose to use fMRI to detect whether a suspect has any familiarity with an individual face. I find that activation in the left inferior frontal gyrus was a reliable discriminator for face familiarity.

## 2 Introduction

The issue of lie detection is critical for law enforcement and the judicial system. The most widespread method currently available is the polygraph test (Meijer *et al*. 2017). In spite of the fact that a 2003 National Academy of Sciences report has concluded that research on the polygraph’s efficacy was inadequate, the government still performs tens of thousands of polygraph tests a year. Some alternative lie detection tools such as voice stress analysis, EEG/Brain fingerprinting, truth drugs, and cognitive chronometry have been tested, but none have exhibited good performance.

New lie detection technology is currently emerging that is based on functional Magnetic Resonance Imaging (fMRI). fMRI measures the haemodynamic response related to neural activity in the brain. It is one of the most recently developed forms of neuroimaging. There has been a recent interest in the use of fMRI for lie detection. Evidence of this is the emergence of several companies: NoLieMRI and Cephose, which were created solely for this purpose. Such companies anticipated that clients such as people facing criminal proceedings and US federal government agencies will seek out their services (Pearson, 2006).

Most lie detection fMRI studies so far have focused on identifying specific regions in the brain that become more active when a person is lying, regardless of what they are lying about (Kozel *et al*., 2005; Langleben *et al*., 2002). Robust group activation have been found that correlated with lying; however, lie prediction based on single subject analysis achieved only 88% accuracy (Langleben *et al*., 2005; Wild, 2005), which is on the same order of accuracy as the polygraph test. The results are also complicated by issues of experimental validity, since lying is cued by the experimenter.

One of the main problems of this general lie detection approach is that it assumes that all acts of deception share common mental operations; however, acts of deception can be very different from one another. So it seems a simplification to even think of deception as a unified class of cognitive mechanisms that can be revealed with a single measurement (Ganis *et al*., 2003, Bles *et al*., 2008) A range of cognitive and motivational processes are involved in deception and thus studies have shown a large variability in areas reported to be involved in lying (Bles *et al*., 2008).

In this work I propose a different fMRI based procedure for testing deception. A way to refine the fMRI lie detection technique is make it problem or domain specific. Hence each lie detection test will be tailored towards a particular cognitive domain. Likewise, the fMRI analysis should be tailored to the specific issue or domain that is being examined. The problem with using a general lie test is that there are too many variables that need to be controlled, such as motivation, attention, emotion, etc. However, in a domain specific test, the control can be tailored to what is being tested. In fMRI tests brain activity is always computed based on certain conditions compared to some control state, and the choice of this control is critical. For example an unfamiliar face is a good control for a familiar face. In this particular work I deal with the issue of face recognition. I propose that a face-specific fMRI-based lie detection system can reveal information about personal familiarity.

In criminal investigations one frequently needs to establish a degree of familiarity between a suspect and a victim, a victim and a witness, or a suspect and a witness. Sometimes this information may be concealed by a witness or suspect for various reasons such as pity, threat, etc. A lie detector that could provide an objective test of familiarity relationship could be very useful in criminal investigations and could be used as circumstantial evidence.

It has been shown in numerous neuroimaging studies that specific regions in the brain are activated during face perception (Ishai *et al*., 2005). The most consistent activation is found in the Fusiform Face Area (FFA), which has been proposed to be a module specialized for face perception (Kanwisher *et al*., 1977). Evidence also exists that suggest that damage to this brain area can be associated with prosopagnosia, a specific face recognition deficit. However, the specific function of this region in processing face stimuli is still under active investigation (Kanwisher, 2000; Haxby *et al*., 2001).

In this work I asked whether the FFA and/or other brain areas are involved in discrimination of familiar faces from unfamiliar faces. Recognition of familiar faces is a complex task, which involves not only retrieval of perceptually coded face images from memory, but also association of semantic and lexical information with those images (Bruce and Young, 1986). Hence I would expect that recognition of familiar face images should lead to much more elaborate activation patterns in the brain than recognition of unfamiliar face images. Previous neuroimaging studies have found various areas that show increased activation to familiar faces when compared to unfamiliar faces (Gorno-Tempini *et al*., 1998; Leveroni *et al*., 2000; Sergent *et al*., 1992; Nakamura *et al*., 2000). This variation is probably due to the large number of mental operations that are involved in recognition of familiar faces. In this study I use a simple block design to examine which brain areas exhibit the most robust activation differences to familiar and unfamiliar faces.Ifind that the left inferior frontal gyrus shows increased activation in all tested subjects when they are viewing famous faces. I believe that this activation is highly robust and can potentially be used to determine if a face is familiar or not in a lie detection test.

## 3 Materials and Methods

### 3.1 Subjects

Ten healthy right-handed volunteers, five males and five females, participated in the experiment (age range 23-35). All participants gave written informed consent, GCO # 04-0361.

### 3.2 Stimuli and Task

This study had two parts. In the first part of the experiment, subjects passively viewed grayscale photographs of unfamiliar faces and houses. This enabled us to anatomically locate any areas which might be specialized for face perception within individual subjects, such as the FFA.

In the second part of the experiment I used a different set of unfamiliar grayscale face photographs and a set of grayscale famous face photographs. Famous and unfamiliar faces were matched for facial expression. All faces were frontal images. Subjects performed a passive viewing task and a gender classification task on two separate runs. In the gender classification task subjects indicated the gender of each stimulus by pressing a button with either their right thumb or left thumb.

In total three time series were obtained from each subject: functional face area localizer, passive viewing and the gender classification task. The functional face area localizer was always performed first and contained 8 blocks, 4 for each stimulus condition. The order of the subsequent two time series was randomized. Each of these latter time series consisted of 12 blocks, 6 for each stimulus condition. All time series began and ended with 16 seconds of rest and contained blocks that were separated by 16 second intervals of rest. The order of the blocks was counterbalanced across time series.

In both part I and Part II stimuli were blocked by condition (famous faces, unfamiliar faces, houses). Each stimulus was presented for 880 milliseconds with 200 millisecond inter-stimulus fixation. Stimuli were presented using E-Prime Software (Psychology Software Tools, Inc.), and were projected by a magnetically shielded LCD video projector onto a translucent screen. The subject viewed the screen through a mirror system.

### 3.3 Imaging

#### 3.3.1 fMRI

fMRI, a noninvasive neuroimaging technique, maps changes in brain haemodynamics that correspond to mental activity (Huettel *et al*., 2004). By using this technique one can map brain structures that participate in specific functions. Increased activity of neurons within a brain region causes an increase in blood flow, thereby increasing the ratio of oxyhemoglobin to deoxyhemoglobin in that region. Brain function is observed by measuring the blood-oxygen-level-dependent (BOLD) response. This measurement is possible because a change in oxygenation causes a change in the magnetic properties of the blood. The fMRI the acquisition is then sensitized to magnetic properties of the underlying tissue.

#### 3.3.2 Protocol

Responses to different images were measured using blood oxygen dependent (BOLD) contrast fMRI with the acquisition of gradient echo-planar (GE-EPI) sequence. Scanning was accomplished at the Mount Sinai Medical Center with a 3.0T Siemens Allegra MRI scanner, equipped with a standard clinical head coil. Each volume was acquired using the following protocol: 32 coronal slices 3mm thick and skip = 1mm, TR=2s, TE=40ms, Flip angle=90 degrees, field of view (FOV)=21cm, matrix=64x64.

## Statistics

Image data were analyzed with statistical parametric mapping software – SPM2 (Welcome Department of Imaging Neuroscience, London, UK www.fil.ion.ucl.ac.uk/spm/). All functional volumes were realigned to the first volume, corrected for motion artifacts, mean adjusted by proportional scaling, normalized onto standard stereotactic space (template provided by Montreal Neurological Institute) and smoothed using an 8mm Gaussian filter in 3D. The time series were high-pass filtered to eliminate low-frequency noise (filter width = 128 seconds). A General Linear Model was used to compute the t-statistic and transform to a unit normalized distribution (Z). Statistical block design analysis comparing the conditions modeled was performed.

The main effect of faces (i.e. the response to faces as compared with the response to houses) and face familiarity (i.e. the response to unfamiliar faces as compared with the response to familiar faces) was analyzed using a linear convolution model with an assumed hemodynamic response function (Friston *et al*., 1995a, 1995b). Clusters were selected that showed a significant effect (P<0.001 uncorrected) with 8 or more contiguous voxels.

## 4 Results

### 4.1 Activation resulting from perception of faces

Two regions were identified that were consistently activated more during perception of faces than houses (P<0.001 with 8 contiguous voxels), as shown in Figures 1 and 2. These areas were the Occipital Face Area (OFA) and the Fusiform Face Area (FFA). Within these regions bilateral activation was found in all subjects, but responses in these “face selective” regions were stronger in the right hemisphere. This was evidenced by the fact that the face responsive clusters in all subjects were larger on the right side than on the left side.

**Figure 1.**
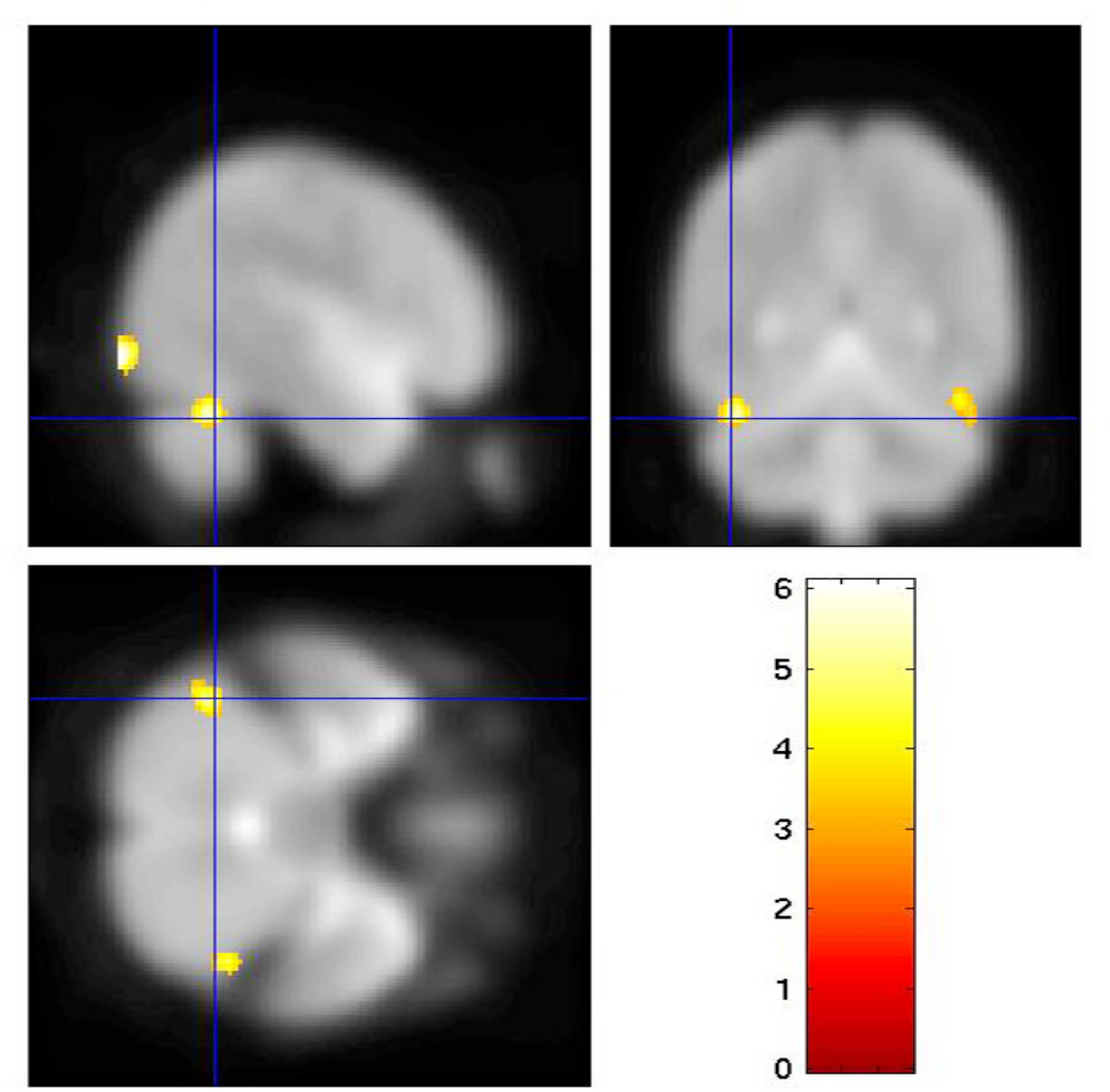
Results of experiment 1, localizer, for one subject. Voxels that were preferentially activated by faces compared to houses are shown. Color scales indicate the degree of statistical significance as measured by a t-test. The Statistical Parametric Map, threshold at P<0.001 uncorrected, is displayed on sagittal, transverse and coronal sections of the fMRI, centered on the maximal voxel in the left fusiform gyrus [x=-44, y=-54, z=-28].

**Figure 2.**
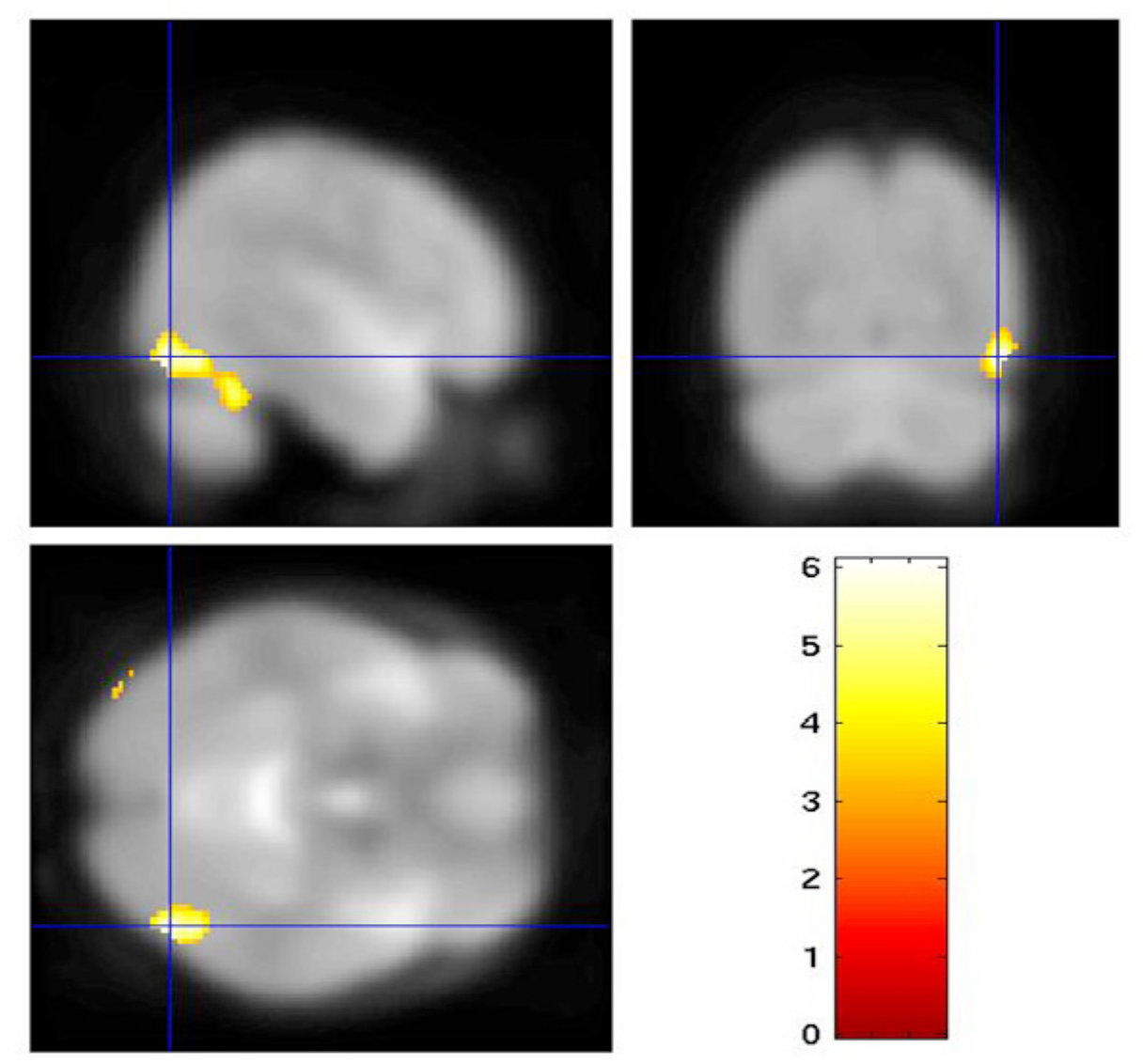
Results of experiment 1, localizer, for one subject. Voxels that were preferentially activated by faces compared to houses are shown. Color scales indicate the degree of statistical significance as measured by a t-test. The Statistical Parametric Map, threshold at P<0.001 uncorrected, is displayed on sagittal, transverse and coronal sections of the fMRI, centered on the maximal voxel in the right occipital gyrus [x=46, y=-74, z=-12].

### 4.2 Activation resulting from perception of famous faces

In the passive viewing task, as shown in Figure 3, the analysis of contrast between familiar and unfamiliar faces revealed a stronger response to familiar faces in the anterior cingulate area and in the left inferior frontal gyrus (LIFG) (P<0.001 with 8 contiguous voxels). Activation in the anterior cingulate is typically associated with attention (Bush *et al*., 1999). Hence I presumed that this activation might result from the fact that familiar faces were more interesting than unfamiliar faces and this caused observers to pay more attention to these stimuli.

**Figure 3.**
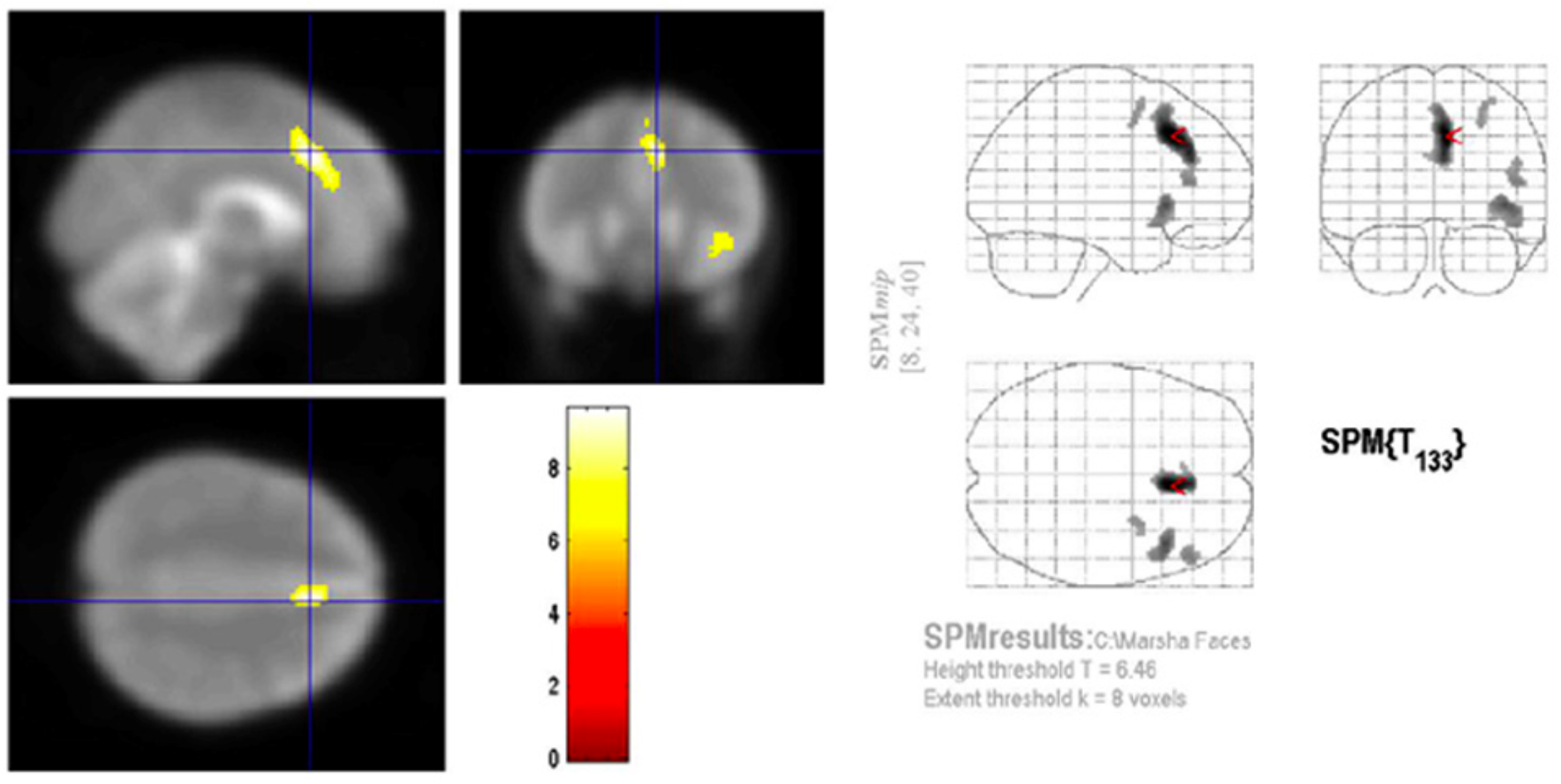
Results of experiment 2 for one subject. Voxels that were preferentially activated by passive viewing of famous faces compared to unfamiliar faces are shown. Color scales indicate the degree of statistical significance as measured by a t-test. The Statistical Parametric Map, threshold at P<0.001 uncorrected, is displayed on sagittal, transverse and coronal sections of the fMRI. The image is centered on the the maximal voxel in the anterior cingulate [x=8, y=24, z=40].

In order to control for attention I asked observers to perform a gender classification task, and, as expected, the activation in the anterior cingulate was eliminated. However, the other activated center, in LIFG, remained active (Figure 4). Activation in LIFG has been implicated in semantic operations, such as generating names of objects belonging to a particular category, such as animals (Costafreda *et al*., 2006). Thus this activation may result from automatic lexical retrieval of famous face names.

**Figure 4.**
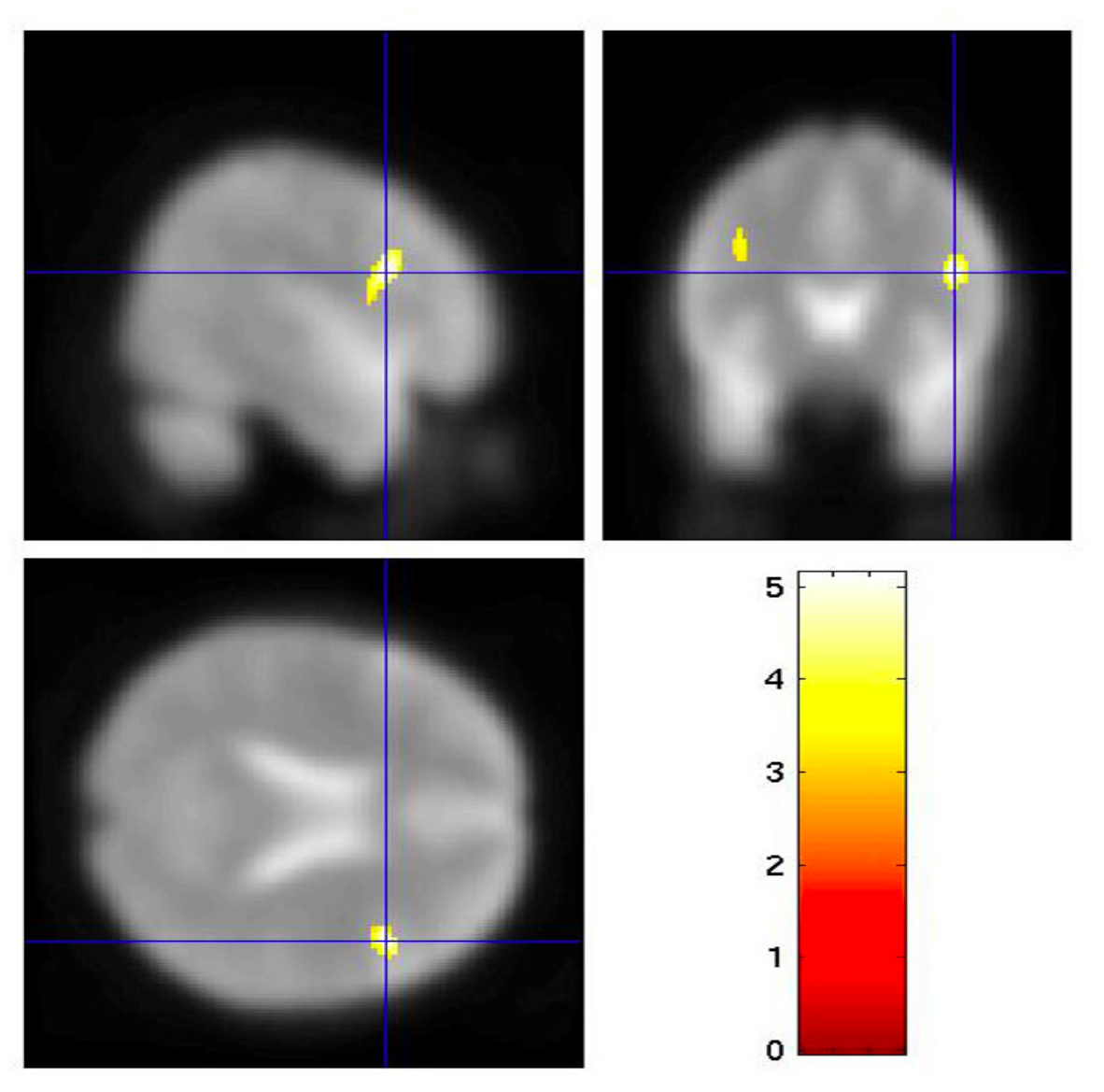
Results of experiment 2 for one subject. Voxels that were preferentially activated by famous faces compared to unfamiliar faces, during the gender classification task, are shown. Color scales indicate the degree of statistical significance as measured by a t-test. The Statistical Parametric Map, threshold at P<0.001 uncorrected, is displayed on sagittal, transverse and coronal sections of the fMRI. The image is centered on the the maximal voxel in the left inferior frontal gyrus [x=46, y=14, z=22].

## 5 Discussion

In this work I propose a lie detection system that is specific to determining one’s familiarity with a face. The neural response evoked by viewing familiar and unfamiliar faces was investigated using fMRI. Robust activation differences were found in LIFG. It is possible that other brain face regions, such as the FFA or OFA, may also be used for performing the familiarity discrimination. In the future, more sensitive data analysis techniques could be used to test this further. More sensitive techniques that are based on pattern classification, such as Support Vector Machines (SVM) and Fisher Linear Discriminant (FLD), are ideal for situations where one must discriminate between two conditions (such as familiar faces and unfamiliar faces). The SVM method is particularly promising because it has been shown that the spatial resolution limitations of fMRI can be overcome by using SVM. This data analysis method has been used successfully to predict the orientation of stimuli based on the fMRI activity in V1, which contains orientation-selective columns only 300-500 μm in width (Kamitani and Tong, 2005).

The principle of lie detection that is proposed for the face recognition domain could potentially be extended into the more general domain of object recognition. In a criminal investigation one may need to establish a connection between an object and a suspect, victim and/or witness. An object could be anything from a weapon to a simple household item that was found at the scene of the crime. To establish a connection an investigator may want to know if someone is familiar with a particular object. fMRI has been used to show that various large clusters of neurons in the ventral temporal cortex (IT) respond selectively to various classes of objects. Category selective regions in IT have been found for cars, birds, chess boards, cats, bottles, scissors, shoes, chairs, and faces (Peissig & Tarr 2007, Reddy & Kanwisher 2006). It is likely that activation patterns in these areas of IT or in other parts of the brain could be used to determine if an object from a particular class is familiar or not and this information could be used for lie detection.

